# Levels of outpatient prescribing for four major antibiotic classes and rates of septicemia hospitalization in adults in different US states

**DOI:** 10.1101/404046

**Authors:** E. Goldstein, S. Olesen, Z. Karaca, C. Steiner, C. Viboud, M. Lipsitch

## Abstract

**Background:** Rates of sepsis/septicemia hospitalization in the US have risen significantly during recent years, and antibiotic resistance and use may contribute to those rates through various mechanisms.

**Methods:** We used multivariable linear regression to relate state-specific rates of outpatient prescribing overall for fluoroquinolones, penicillins, macrolides, and cephalosporins between 2011-2012 to state-specific rates of hospitalization with septicemia (ICD-9 codes 038.xx present anywhere on discharge diagnosis) in each of the following age groups of adults: (18-49y, 50-64y, 65-74y, 75-84y, 85+y) reported to the Healthcare Cost and Utilization Project (HCUP) between 2011-2012, adjusting for additional covariates, and random effects associated with the ten US Health and Human Services (HHS) regions.

**Results:** Rates of penicillin prescribing were positively associated with septicemia hospitalization rates in the analyses for persons aged 50-64y, 65-74y, and 74-84y. Percent African Americans in a given age group was positively associated with septicemia hospitalization rates in the analyses for persons aged 75-84y and over 85y. Average minimal daily temperature was positively associated with septicemia hospitalization rates in the analyses for persons aged 18-49y, 50-64y, 75-84y and over 85y.

**Conclusions:** Our results suggest positive associations between the rates of prescribing for penicillins and the rates of sepsis hospitalization in US adults aged 50-84y. Further studies are needed to understand the potential effect of antibiotic replacement in the treatment of various syndromes, such as replacement of fluoroquinolones by other antibiotics, possibly penicillins following the recent US FDA guidelines on restriction of fluoroquinolone use, on the rates of sepsis hospitalization.

## Introduction

Rates of hospitalization with septicemia, sepsis, associated mortality and monetary costs have been rising rapidly during the past decades in the US [1-4]. While changes in diagnostic practices contributed to the rise in the rates of hospitalization with septicemia/sepsis in the diagnosis [5,6], those changes cannot fully explain that rise in hospitalization rates, particularly prior to 2010 [7]. Indeed, trends in the rates of hospitalization with any diagnosis of sepsis between 2003-2009 closely resembled the trends in the rates of hospitalizations that involved infection and the use of mechanical ventilation (Figure 1 in [7]). Antibiotic resistance and use may also contribute to septicemia/sepsis hospitalization rates [8]. The relation between infections with antibiotic-resistant bacteria and survival for sepsis, including the effect of initially appropriate antibiotic therapy (IIAT) is suggested by a number of studies, e.g. [9,10]. However, less is known about the relation between levels of antibiotic use, as well as prevalence of antibiotic resistance and rates of hospitalization with septicemia/sepsis. Antibiotic resistance can facilitate the progression to severe disease when infections not cleared by antibiotics prescribed during outpatient and/or the inpatient treatment devolve into a more severe state, including sepsis. For example, antibiotic resistance in Enterobacteriaceae, including fluoroquinolone resistance in *Escherichia coli* was found to be associated with a more severe presentation in urinary tract infections (UTIs) [11,12], while prevalence of fluoroquinolone resistance in *E. coli* was strongly correlated with rates of septicemia hospitalization in different US states for adults aged over 50y [8]. In a study of cases of *E. coli* bacteremia in England, urogenital infection had been treated in 310/891 (34.8%) cases in the 4 weeks preceding bacteremia and this sub-population differed very significantly in its antibiotic resistances [13], suggesting treatment failure due to presence of antibiotic resistance prior to the onset of bacteremia. Antibiotic use is one important driver of the prevalence of antibiotic resistance [14-19] and thus may contribute to sepsis incidence. Antibiotic use has been implicated as a cause of resistance not only to the drug class used, but to other drugs. For example, fluoroquinolone use was found to be associated with methicillin-resistant *S. aureus* (MRSA) infections [20-23], with some of those infections leading to sepsis outcomes, while amoxicillin use was found to be associated with trimethoprim resistance in Enterobacteriaceae [14], with trimethoprim-resistant urinary tract infections (UTIs) leading to bacteremia outcomes [15]. While causality is not conclusively proven in these studies, it is plausible given the fact that resistance to different drug classes tends to cluster in bacterial populations, leading to the phenomenon of co-selection [24,25].

Earlier work has shown a relationship, including spatial correlations between the rates of antibiotic consumption and prevalence of antibiotic resistance, e.g. [14-19]. At the same time, there is limited information on the relationship between levels of prescribing for different antibiotic types/classes, and rates of sepsis hospitalization. Moreover, that relationship may be context-specific and affected by various factors such as the use of other antimicrobials and prevalence of cross-resistance [26,27], local patterns of transmission/acquisition of antibiotic-resistant infections, demographic and climatic differences, etc. This study examines the relationship between the rates of prescribing for different antibiotic classes and rates of hospitalization for septicemia/sepsis for population-level data in the US. We use a regression framework to relate the state-specific rates of outpatient prescribing of fluoroquinolones, penicillins, macrolides, and cephalosporins in the US CDC Antibiotic Patient Safety Atlas data [28] to state-specific rates of hospitalization with septicemia (ICD-9 codes 038.xx present on the discharge diagnosis) in different age groups of adults recorded in the Healthcare Cost and Utilization Project (HCUP) data [29], adjusting for additional factors that may affect septicemia hospitalization rates. The framework for the analyses in this paper is similar to the framework for the analyses in our parallel study on the relation between antibiotic prescribing and septicemia mortality [30]. We hope that such analyses can lead to further study of the contribution of different antibiotics to the rates of sepsis hospitalization, including the utility of antibiotic stewardship and shift in prescribing from some antibiotics to others in the treatment of certain syndromes for reducing the levels of sepsis hospitalization in the US.

## Methods

### Data

We used data between 2011-2012 on counts of hospitalization with a septicemia diagnosis (both primary and contributing, [*ICD-9*] codes 038.xx on the discharge diagnosis) from the State Inpatient Databases of the Healthcare Cost and Utilization Project (HCUP), maintained by the Agency for Healthcare Research and Quality (AHRQ) through an active collaboration [29]. We will henceforth call these hospitalizations septicemia hospitalizations, even though some of them may involve low levels of bloodstream bacterial infection. The database [29] contains hospital discharges from community hospitals in participating states. Forty-two states reported septicemia hospitalization data between 2011-2012 for each of the five adult age groups included in our analyses: 18-49y, 50-64y, 65-74y, 75-84y, 85+y. Those states are Alaska, Arkansas, Arizona, California, Colorado, Connecticut, Florida, Georgia, Hawaii, Iowa, Illinois, Indiana, Kansas, Kentucky, Louisiana, Massachusetts, Maryland, Minnesota, Missouri, Montana, North Carolina, North Dakota, Nebraska, New Jersey, New Mexico, Nevada, New York, Ohio, Oklahoma, Oregon, Rhode Island, South Carolina, South Dakota, Tennessee, Texas, Utah, Virginia, Vermont, Washington, Wisconsin, West Virginia, and Wyoming. Aggregate state-level hospitalization data were used in the analyses, with no informed consent from the participants sought.

We extracted data on the annual state-specific per capita rates of outpatient prescribing for four classes of antibiotics: fluoroquinolones, penicillins, macrolides, and cephalosporins in 2011 and 2012 from the US CDC Antibiotic Patient Safety Atlas [28]. Average annual state-specific outpatient prescribing rates (per 1,000 residents) for each class of antibiotics between 2011-2012 were then estimated as the average for the corresponding rates in 2011 and 2012.

Annual state-specific population estimates in each age group of adults were obtained as the yearly July 1 population estimates in [31]. Data on median household income for US states between 2011-2012 were extracted from [32]. Data on average minimal daily temperature for US states were obtained from [33]. Data on the percent of state residents in different age groups who lacked health insurance were extracted form the US Census Bureau database [34].

### Regression model

For each age group of adults, (18-49y, 50-64y, 65-74y, 75-84y, 85+y), we applied multivariable linear regression to relate the state-specific outpatient prescribing rates (per 1,000 state residents) for fluoroquinolones, penicillins, macrolides, and cephalosporins between 2011-2012 [28] (covariates) to the state-specific rate of septicemia hospitalization in the given age group between 2011 and 2012 [29] (dependent variable). Besides the antibiotic prescribing rates, the other covariates used in the model were the state-specific median household income, the state-specific average minimal daily temperature, and percentages of state residents in a given age group who were African American, as well as those who lacked health insurance (for age groups under 65 years). We note that temperature may influence bacterial growth rates and/or transmission mediated effects [35,36], which in turn may affect both the prevalence of antibiotic resistance and the acquisition/severity of bacterial infections. We also note that African Americans have elevated rates of sepsis compared to other races [37]. To adjust for additional factors not accounted for by the covariates used in the model, we include random effects for the ten Health and Human Services (HHS) regions in the US. Specifically, for each state *s* that reported the corresponding septicemia hospitalization data between 2011-2012 in [29], let *HR*(*s*) be the state-specific rate of hospitalizations with septicemia in the given age group per 10,000 residents, *A*_*i*_(*s*) (*i = 1, ‥, 4*) be the state-specific average annual outpatient prescribing rates between 2011-2012, per 1,000 state residents, for the four studied classes of antibiotics; *I*(*S*) be the median state-specific household income (in $1,000s) between 2011-2012; *MT*(*s*) be the average minimal daily temperature (°F) between 2005-2011 for the state; *AA*(*s*) be the percent of African Americans among the state residents in a given age group between 2011-2012; *LHI*(*s*) be the average annual age-specific percent of state residents who lacked health insurance between 2011-2012 (for non-elderly age groups); and *RE*(*s*) be the random effect for the corresponding HHS region. Then:

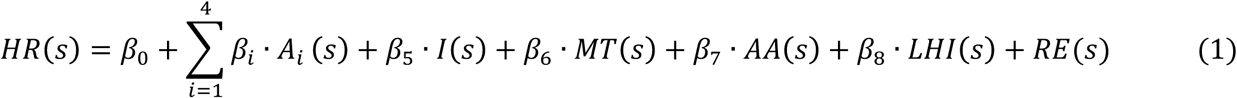

This regression was performed five separate times, once for the outcome in each age group, with the health insurance and percent African-American covariates being age-group-specific, and the others being the same between regressions. The random effect was estimated independently in each regression.

## Results

State-specific rates of outpatient prescribing of *all* antibiotics per 1,000 residents in [28] are correlated with the state-specific rates of septicemia hospitalization in each age group of adults in [29] (cor = 0.44 (95% CI (0.15,0.65)) for persons aged 18-49y, 0.51 (0.24,0.70) for persons aged 50-64y, 0.49 (0.21,0.69) for persons aged 65-74y, 0.41 (0.12,0.63) for persons aged 75-84y, and 0.3 (-0.01,0.55) for persons aged 85+y).

Partitioning the relationship between the rates of antibiotic prescribing and rates of septicemia hospitalization by antibiotic class is challenging, in part because of the high positive correlations between the rates of outpatient prescribing for different antimicrobial classes at the state level. Table 1 presents the correlations between the state-specific rates of outpatient prescribing for different classes of antibiotics (fluoroquinolones, penicillins, macrolides, and cephalosporins) in [28]. All the correlations in Table 1 are high, with point estimates above 0.76 for each pair of classes of antibiotics.

**Table 1:** Correlations between the state-specific rates of outpatient prescribing of fluoroquinolones, penicillins, macrolides, and cephalosporins in [28] between 2011-2012.

|  | Fluoroquinolones | Penicillins | Macrolides | Cephalosporins |
| --- | --- | --- | --- | --- |
| Fluoroquinolones | 1 |  |  |  |
| Penicillins | 0.8<br>(0.66,0.89) | 1 |  |  |
| Macrolides | 0.88<br>(0.79,0.94) | 0.83<br>(0.7,0.91) | 1 |  |
| Cephalosporins | 0.76<br>(0.59,0.86) | 0.82<br>(0.69,0.9) | 0.82<br>(0.68,0.9) | 1 |

Table 2 presents the estimates (regression coefficients) for the multivariable regression model (eq. 1) that relates the state-specific rates of outpatient prescribing for the four studied classes of antibiotics, the median household income, average minimal daily temperature, and percentages of state residents in a given age groups who were African American, as well as those who lacked health insurance (for non-elderly age groups) to the state-specific septicemia hospitalization rates in different age groups of adults between 2011-2012 [29]. The regression coefficients for the different antibiotic classes estimate the change in septicemia hospitalization rates (per 10,000 individuals in the given age group) when the annual rate of outpatient prescribing of the corresponding antibiotic class (per 1,000 residents) increases by 1. The regression coefficients were positive for the rates of prescribing of penicillins in the analyses for persons aged 50-64y, 65-74y, and 75-84y. The regression coefficients for the average minimal daily temperature were positive in the analyses for persons aged 18-49y, 50-64y, 75-84y and over 85y. Percent African Americans in a given age group was positively associated with septicemia hospitalization rates for persons aged 75-84y and over 85y.

**Table 2:** Regression coefficients in eq. 1 for the different covariates in the model given by eq. 1 for different age groups. The coefficients for the different antibiotic classes estimate the change in the annual septicemia hospitalization rates (per 10,000 individuals in a given age group) when the annual rate of outpatient prescribing of the corresponding antibiotic class (per 1,000 residents) increases by 1. ND=not done because persons >64 years old are eligible for Medicare.

|  | Aged<br>18-49y | Aged<br>50-64y | Aged<br>65-74y | Aged<br>75-84y | Aged<br>85+y |
| --- | --- | --- | --- | --- | --- |
| Fluoroquinolones<br>(prescription per<br>1000 residents/y) | 0.08<br>(-0.07,0.22) | 0.18<br>(-0.15,0.51) | 0.36<br>(-0.3,1) | 0.7<br>(-0.45,1.9) | 1.3<br>(-1.1,3.7) |
| Penicillins<br>(prescription per<br>1000 residents/y) | 0.04<br>(-0.03,0.12) | **0.19**<br>**(0.02,0.37)** | **0.48**<br>**(0.12,0.84)** | **0.81**<br>**(0.17,1.4)** | 1.2<br>(-0.14,2.5) |
| Cephalosporins<br>(prescription per<br>1000 residents/y) | -0.02<br>(-0.09,0.05) | -0.03<br>(-0.19,0.13) | -0.08<br>(-0.42,0.25) | -0.28<br>(-0.88,0.31) | -0.69<br>(-1.9,0.51) |
| Macrolides<br>(prescription per<br>1000 residents/y) | -0.03<br>(-0.11,0.04) | -0.12<br>(-0.29,0.04) | -0.26<br>(-0.6,0.08) | -0.39<br>(-0.99,0.22) | -0.55<br>(-1.8,0.7) |
| Median household<br>income (\$1000) | -0.09<br>(-0.29,0.1) | -0.17<br>(-0.58,0.24) | 0.31<br>(-0.5,1.1) | 1.2<br>(-0.24,2.6) | **3.3**<br>**(0.42,6.2)** |
| Average minimal daily<br>temperature (°F) | **0.2**<br>**(0,0.39)** | **0.57**<br>**(0.12,1)** | 0.78<br>(-0.14,1.7) | **1.6**<br>**(0.03,3.2)** | **3.9**<br>**(0.59,7.3)** |
| Percent African<br>Americans | -0.05<br>(-0.22,0.12) | 0.15<br>(-0.28,0.59) | 0.79<br>(-0.23,1.8) | **2.3**<br>**(0.32,4.2)** | **5.3**<br>**(1.1,9.5)** |
| Percent lacking health<br>insurance | 0<br>(-0.25,0.24) | -0.16<br>(-0.91,0.59) | ND | ND | ND |

## Discussion

While antimicrobial use can contribute to the rates of hospitalization with septicemia and sepsis (see the Introduction), the strength of the relation between antibiotic prescribing rates and rates of septicemia/sepsis hospitalization may vary with antibiotic types/classes and depend on a number of factors including patterns of resistance to different antibiotics. In this study, we use multivariable regression to examine the relation between the rates of antibiotic prescribing and the rates of septicemia hospitalization in the US context. Our analyses suggest the association between the state-level rates of outpatient prescribing of penicillins and rates of septicemia hospitalization in persons aged 50-64 years, 65-74 years, and 75-84 years. We note that our companion paper [30] has shown an association between the state-level rates of outpatient prescribing of penicillins and rates of septicemia mortality in older adults. We also note that earlier work has shown a relationship, including spatial correlations between the rates of antibiotic prescribing and antibiotic resistance, e.g. [14-19]. The findings in this paper, of our earlier studies [19,30], as well as the high prevalence of resistance to penicillins in both the Gram-negative and Gram-positive bacteria (e.g. [27,38,39]) support the notion that the use of penicillins in the US is associated with prevalence of antibiotic resistance, while the combination of antibiotic use (particularly for penicillins) and antibiotic resistance is associated with the rates of sepsis hospitalization and mortality in older US adults. Further studies of various related phenomena, including patterns of resistance to different antibiotics for different bacterial infections are needed to better understand the potential effect of antibiotic, particularly penicillin replacement in the treatment of various syndromes on the rates of sepsis hospitalization and mortality. Additionally, we have found that prevalence of African Americans among older adults is associated with rates of septicemia hospitalization, which agrees with earlier findings that rates of sepsis hospitalization among African Americans are elevated [37]. Our work also suggests that average minimal daily temperature is positively associated with the rates of septicemia hospitalization in four out of five age groups of adults, which resonates with earlier findings about the association between temperature and antibiotic resistance [35,36,19].

Patterns of resistance to different antibiotics modulate the relation between antibiotic prescribing and rates of sepsis hospitalization. For example, in the UK, outpatient amoxicillin prescribing rates are high ([40], Figure 3.4), and amoxicillin use is associated with the prevalence of trimethoprim resistance [14], while trimethoprim use in turn is associated with the rates of UTI-related *E. coli* bacteremia [15]. The latter association is presumably mediated by the treatment of trimethoprim-resistant UTIs with trimethoprim. Another antibiotic relevant to the growth in the rates of *E. coli*-associated bacteremia in England is amoxicillin-clavulanate (co-amoxiclav). Levels of *E. coli* and *Klebsiella*-associated bacteremia were continuing to rise in England after 2006 while reduction in fluoroquinolone and cephalosporin use was taking place [18,41]. Amoxicillin-clavulanate prescribing in England increased significantly between 2006-2011 [42], and incidence of bacteremia with *E. coli* strains resistant to co-amoxiclav began to increase rapidly after 2006 ([43], Figure 4), with co-amoxiclav resistance in *E. coli* bacteremia exceeding 40% in 2014 [40]. Those findings from the UK suggest that data on resistance patterns for different antibiotics in different bacteria contributing to various syndromes should be considered when new antibiotic prescribing practices (such as the increase in the use of penicillins in England accompanying the decrease in the use of fluoroquinolones and cephalosporins) are being phased in. Recently, US FDA has recommended the restriction of fluoroquinolone use for certain conditions (such as uncomplicated UTIs) due to potential adverse effects [44]. Those recommendations may have benefits in terms of reducing the rates of certain severe bacterial infections, e.g. invasive MRSA infections, and *C. difficile* infections, as also suggested by the UK experience in reduction of fluoroquinolone and cephalosporin prescribing [45,46,20-23]. At the same time, the uptake, as well as the public health benefit of recommendations to reduce one antibiotic class [44] may be enhanced by specific recommendations about choice of alternative treatments. For example, the UK recently updated prescribing guidelines for UTIs, with nitrofurantoin generally recommended as the first-line option [47]. We also note that our results, as well as the UK experience suggest that replacing fluoroquinolones by penicillins in older adults might not be optimal for reducing the rates of severe bacterial infections, including sepsis, as well as the rates of associated mortality [30].

Our analyses have found suggestive evidence of an association between the rates of fluoroquinolone prescribing and rates of sepsis hospitalization, but in no age group did the confidence bounds for this association exclude zero. Cephalosporin and macrolide prescribing, on the other hand, had suggestive negative associations, but again all confidence intervals included zero. The strength and clarity of positive associations for penicillin might partly be related to the fact that the use of certain other antibiotics, particularly penicillins in the treatment of various syndromes may be associated with higher rates of sepsis hospitalization as prevalence of resistance to penicillins in infections with both Gram negative and Gram positive bacteria is high, e.g. [27,38,39]. If (like all models) this model is somewhat mis-specified, competition with antibiotics that are more likely to promote sepsis outcomes could bias downward the strength of the association between the use of certain antibiotics and the rates of sepsis hospitalization. Additionally, various sources of noise, such as the fact that we related antibiotic prescribing rates in the population overall to rates of septicemia hospitalization in different age groups, might have reduced the precision of our estimates. We note the high prevalence of resistance to fluoroquinolones for certain syndromes that contribute to the volume of septicemia hospitalizations, like urinary tract infections (UTIs) [27,26]. Additionally, prevalence of cephalosporin resistance and the frequency of extended-spectrum beta-lactamase (ESBL) production, including in Gram-negative bacteria is growing [48]. Macrolide use could potentially contribute to the rates of sepsis as macrolides are commonly prescribed for the treatment of certain syndromes that are major causes of sepsis, notably respiratory diseases, both chronic [49] and acute, including pneumonia [50]. On the other hand, macrolides are used relatively infrequently in the treatment of urinary tract and gastrointestinal infections. A UK study found that for a high proportion of Gram-negative bacteremia, the main foci of infection were either urinary tract or abdomen/biliary tract [51]. Overall, the potential benefits of antibiotic stewardship for penicillins may be the most important findings of this paper, while options for antibiotic replacement require further investigation.

Our study has some additional limitations. The study is ecological, but for exposures such as antibiotic use, this may be the best design because effects of antibiotic use on resistance go beyond the treated individual to those with whom transmission (directly or indirectly is possible). The antibiotic-sepsis incidence associations we find estimate causal effects only if the model is well-specified and all confounders are accounted for in the analysis. We have included several relevant demographic and geographic variables – income and average minimal daily temperature, percentages of residents who are African American, and those who lack health insurance – with known or hypothesized associations with both antibiotic use and sepsis. We have used age stratification in the outcome to deal with confounding by age composition.

Nevertheless, there may be misspecification and residual confounding (such as geographic differences in coding practices for septicemia). Studies using other methods and in other populations are needed to test the consistency of our findings. To adjust for potential effects of confounding/model misspecification, we have included random effects for the ten US Health and Human Services regions, which led to improvements in model fits. We also note that reverse causality associated with antibiotic treatment for sepsis is unlikely to have a significant contribution to our results because we are considering the relation between *outpatient* prescribing rates and septicemia hospitalizations. While the HCUP data utilized in our study generally cover about 97% of all community hospitalizations in the US [29], state-specific variability in the proportion of septicemia hospitalizations that are covered by the HCUP data is possible. Diagnostic practices for septicemia vary by state. It is uncertain whether states with higher antibiotic prescribing rates have more inclusive criteria for a septicemia diagnosis (boosting the association between antibiotic prescribing rates and septicemia hospitalization rates), or vice versa. For example, California has relatively low antibiotic prescribing rates [52] and apparently more inclusive criteria for a septicemia/sepsis diagnosis that translate into lower case fatality rates for such hospitalizations compared to the national average, e.g. [53]. On the other hand, some of the states with low antibiotic prescribing rates, possibly outside the Northeastern US, may have more strict criteria for a septicemia/sepsis diagnosis compared to the national average. Finally, data on outpatient antibiotic prescribing in the whole population [28] were used in the regression model for which the outcomes were age-specific rates of septicemia hospitalization [29]. We expect that those sources of noise/incompatibility should generally reduce precision and bias toward null results rather than creating spurious associations.

We believe that despite those limitations, our findings indicate a possible causal association between the use of penicillins, and the rates of sepsis hospitalization in US adults aged between 50-84y, suggesting the potential benefits of antibiotic stewardship. We hope that this ecological study would lead to further investigations of the relation between antibiotic prescribing practices and sepsis hospitalizations in different contexts, including the potential effect of replacing penicillins by other antibiotics in the treatment of certain syndromes on the rates of sepsis hospitalization. Finally, we believe that a comprehensive, long-term approach for controlling the rates of severe outcomes associated with bacterial infections, including sepsis should include not only the adoption of appropriate antibiotic prescribing practices but also the introduction of new antibiotics [54].

## Acknowledgement

We would like to thank the HCUP Partner states that voluntarily provide their data to the project, and without whom this research would not be possible (https://www.hcup-us.ahrq.gov/partners.jsp). This study does not necessarily represent the views of the NIH or the US Government.

